# Early-life feeding accelerates gut microbiome maturation in piglets

**DOI:** 10.1101/2020.09.30.320275

**Authors:** R. Choudhury, A. Middelkoop, W.J.J. Gerrits, B. Kemp, J.E. Bolhuis, M. Kleerebezem

**Affiliations:** Host-Microbe Interactomics Group, Department of Animal Sciences, Wageningen University & Research, P.O. Box 338, 6700 AH Wageningen, The Netherlands; Adaptation Physiology Group, Department of Animal Sciences, Wageningen University & Research, P.O. Box 338, 6700 AH Wageningen, The Netherlands; Animal Nutrition Group, Department of Animal Sciences, Wageningen University & Research, P.O. Box 338, 6700 AH Wageningen, The Netherlands

**Keywords:** early-life, development, fibre, feeding strategies, microbiota, neonatal, pig, suckling, weaning

## Abstract

Early-life microbiome perturbations have been suggested to have important effects on host development, physiology, and behaviour, which can persist throughout life. We hypothesise that early feeding (access to a pre-weaning fibrous diet) can affect gut microbiome colonisation and development in neonatal piglets. In this longitudinal study, a customised fibrous diet was provided to early-fed piglets (EF; 6 litters) starting two days after birth until weaning (28 days of age) in addition to mother’s milk, whereas control piglets (CON; 4 litters) suckled sow’s milk only. Rectal swabs were collected at multiple timepoints until six weeks of age (i.e., 2 weeks post-weaning) to investigate intestinal microbiota composition development over time using 16S rRNA gene profiling (n = 10 piglets per treatment). We observed a dynamic intestinal microbiota colonisation pattern during the pre-weaning period in both treatment groups, which rapidly stabilised within 2 weeks post-weaning. The microbial (alpha) diversity increased with age and seemed to reach a plateau in the early post-weaning time-point (day+5). The homogenous post-weaning microbiota was represented by microbial groups including *Prevotella, Roseburia, Faecalibacterium, Ruminococcus, Megasphaera, Catenibacterium* and *Subdoligranulum*. Remarkably, early feeding of neonatal piglets resulted in accelerated maturation of the intestinal microbiota at pre-weaning time-points, characterised by increased rate of microbial diversity and expanded colonisation of typical post-weaning associated microbial groups (mentioned above) at pre-weaning stages. The acceleration in EF piglets was illustrated by the simultaneous emergence of typical post-weaning-associated microbial groups and a more rapid decline of typical early-life/pre-weaning microbial genera. In addition, the individual eating behaviour scores of the piglets quantitatively correlated with the accelerated change of their microbiome. Overall, these findings show the importance of early-life nutritional strategies to influence the gut microbiota maturation in piglets.

## INTRODUCTION

The mammalian gastrointestinal tract (GIT) is inhabited by complex and diverse microbial communities that influence host health and disease. Rapid advances in next generation sequencing have facilitated the understanding of the factors that shape this complex community. The complex ecosystem development initiates by microbial groups starting to assemble shortly after birth (derived from the mother and environment) and gradually stabilises over time (Schokker et al., 2015; Timmerman et al., 2017). In early-life, the influence of colonising gut microbiota on intestinal development is crucial as in this period, the microbiota is considered essential for appropriate development and programming of the mucosal immune response (Schokker et al., 2015; Zhuang et al., 2019). Based on the plasticity of the mucosal system during this early-life development phase, it has been proposed as a “window of opportunity” where perturbations may have long-lasting impacts on health and welfare (Putignani et al., 2014; Nowland et al., 2019).

The transition from mother’s milk to solid food, commonly known as weaning, is normally a gradual process in young mammals. In nature, the weaning process of piglets approaches completion between 17 - 20 weeks of age approximately (Newberry and Wood-Gush, 1985; Jensen and Stangel, 1992). However, in modern swine industry practice, piglets are weaned abruptly around 3 - 4 weeks of age, a time-period that coincides with the developmental changes of the gastrointestinal system (Moeser et al., 2017). Evidently, early weaning is a highly dynamic and stressful event as the piglets deal with sudden changes in diet and environment, including the separation from their mother and littermates. Weaning stress is typically associated with low feed intake, sub-optimal weight gain, diarrhoea episodes and maladaptive behaviour, leading to compromised animal health and welfare, increased piglet mortality and economic losses (Le Dividich and Sève, 2000; Bruininx et al., 2002; Heo et al., 2013; Everaert et al., 2017; Gresse et al., 2017; Pluske et al., 2018). Gut microbiota dysbiosis (characterised by microbial imbalance, intestinal inflammation, reduced gut barrier function, and increased abundance of potential pathogens) is one of the factors that could contribute to the weaning transition problems, and is considered as one of the leading causes of post-weaning diarrhoea and associated GIT infections in piglets (Gresse et al., 2017; Pluske et al., 2018). The two major pathogens, identified as causative agents in post-weaning diarrhoea of piglets, are *Salmonella enterica serovar Typhimurium* and especially enterotoxigenic *Escherichia coli* (ETEC), causing high pig mortality rates worldwide (Gresse et al., 2017; Tran et al., 2018). Although weaning transition has been associated with intestinal dysbiosis, the role of the microbiota changes in the overall health-risks at weaning remains elusive. This highlights the need for further research to decipher the impact of early-life perturbations on gut microbiota colonisation and its consequences during the weaning transition as well as later life in pigs.

In neonatal piglets, the microbiota colonisation is a dynamic process with rapid microbial shifts from the initial pioneering microbial groups (present during the first weeks of life) to the changing microbial populations eventually reach a climax, adult-like microbial community (Guevarra et al., 2019). Several studies (Kim et al., 2011; Alain et al., 2014; Frese et al., 2015; Mach et al., 2015; Niu et al., 2015; Slifierz et al., 2015; Zhao et al., 2015; Chen et al., 2017; Ke et al., 2019; Wang et al., 2019) have evaluated the microbiota development in pigs, analysing the age-related microbial shifts, employing either faecal samples or rectal swabs. For instance, Chen and colleagues (Chen et al., 2017) characterised longitudinal changes in faecal microbiota of suckling piglets at four different ages and described a quickly shifting intestinal microbiota around weaning that stabilised in a period of approximately 10 days post-weaning, with microbial groups like *Bacteroides, Escherichia/Shigella* group enriched during pre-weaning ages, and *Prevotella* predominating the post-weaning ages where it accumulated to approximately 25% of the overall gut microbiota community.

It is well established that diet can be considered a major factor that can shape the intestinal microbiota in mammals. Dietary fibres, abundant in common plant-based feedstuffs, pass through the small intestine in an undigested form and are fermented in the distal ileum and colon, stimulating the growth of microbes. Microbial fermentation of undigested fibres usually takes place in the distal part of the gastrointestinal tract, resulting in the formation of short-chain fatty acids (SCFA), that are known to influence physiological functioning of the intestines such as formation and protection of intestinal barrier as well as host defence and inflammatory responses (Den Besten et al., 2013; Furusawa et al., 2013; van der Beek et al., 2017; Xiong et al., 2019). Several studies have evaluated the impact of dietary fibres on intestinal microbiota composition, focussing mostly on weaned or growing pigs (Yao, 2008; Ivarsson et al., 2012; Liu et al., 2012, 2018; Haenen et al., 2013; Dicksved et al., 2015; Umu et al., 2015, 2018; Burbach et al., 2017; Zhao et al., 2018; Soler et al., 2018; Yin et al., 2019; Chen et al., 2020). However, only a handful of studies have modulated the early-life “window of opportunity” using dietary treatments in neonatal piglets and evaluated the impact of fibres on microbiota composition (Shim et al., 2005; Berding et al., 2016; Zhang et al., 2016; Mu et al., 2017; Schokker et al., 2018; Van Hees et al., 2019). Most of these studies assessed the effect of specific dietary fibres on the intestinal microbiota at a single time-point, and employed outdated methods of analysis like microbiota cultivation, 16S DNA-microarray technology, and/or qPCR analysis of selected microbial groups. Overall, they report alteration in microbiota composition due to the pre-weaning dietary intervention, especially focussed on increased abundances of genera like *Lactobacillus and Bifidobacterium*, and/or decreased abundances of potentially pathogenic species like *Escherichia coli, Streptococcus suis* or *Clostridium perfringens*.

In the present study, we provided a customised fibrous diet to a group of suckling piglets from two days of age, intending to familiarise them with the consumption of solid feed and to investigate the impact on their gut microbiota compared to control piglets that did not receive pre-weaning solid feed. Both groups of piglets revealed substantial dynamics of the gut microbiome during the pre-weaning period, characterized by the appearance and disappearance of specific microbial genera over time. Additionally, our findings establish that pre-weaning fibrous feed can accelerate gut microbiome colonisation towards a more ‘mature’ microbiome, that resembles that of post-weaning microbial composition. Moreover, the magnitude of this acceleration effect appeared to be quantitatively related to the amount of pre-weaning feed intake as deduced from observational studies. Finally, the pre-weaning provision of fibrous feed reduced variability of post-weaning weight gain, and resulted in a trend towards higher body weight shortly after weaning.

## MATERIALS AND METHODS

### Animals and experimental design

The Animal Care and Use committee of Wageningen University & Research (Wageningen, The Netherlands) approved the protocol of the experiment (AVD104002016515). The protocol is in accordance with the Dutch law on animal experimentation, which complies with the European Directive 2010/63/EU on the protection of animals used for scientific purposes. Ten multiparous Topigs-20 sows (parity 3-5) housed and inseminated at research facility Carus (Wageningen University & Research, The Netherlands), were divided into two groups (n=6 litters for early-fed or EF group; and n=4 litters for control or CON group) based on sow’s parity, body weight and genetic background. Within two days after birth, the litter size was set to maximum 14 piglets per litter (Tempo x Topigs-20) with no cross-fostering. The new-born piglets were cohoused with sow and littermates till weaning (28 days of age) and received ear tags for individual identification and an iron injection, standard to pig husbandry practice. From two days of age, the neonatal piglets belonging to the EF group were given access to customised fibrous feed (**Supplementary Table 1**) *ad libitum* in addition to suckling sow’s milk whereas the CON group suckled sow’s milk only. Briefly, the feed contained 26% dietary non-starch polysaccharides, mainly originating from sugar beet pulp (4%), oat hulls (4%), inulin (4%), galacto-oligosaccharides (5%) and high amylose maize starch (4%) as fibrous ingredients. A subset of piglets (n=64; 32 for each group, EF and CON) were weaned at four weeks of age and followed for two weeks post-weaning. At weaning, piglets were mixed within the same treatment group and housed in separate pens with four unfamiliar piglets per pen (i.e., 4 piglets per pen originating from different litters within treatment). After weaning, all piglets had *ad libitum* access to commercial weaner diet (Inno Speen Pro, Coppens Diervoeding, Helmond, The Netherlands). Additional information about the housing and management have been described in detail in a previous study (Middelkoop et al.).

### Eating behaviour by video observation

The eating behaviour of piglets was assessed by video recordings from two days of age till weaning. For recognition during behavioural observations, piglets were individually numbered on their back using dark permanent hair dye. Eating frequency of individual EF piglets was determined daily from 07:00 to 19:00 hours via video observations as an estimate for pre-weaning solid feed intake. From the video observations, the amount of time spent eating or “eating time” was evaluated. When an EF piglet placed its snout into the trough for a minimum of 5 seconds (s), the behaviour was scored as eating (Pajor et al., 1991; Adeleye et al., 2014). The eating time was categorised into short (5 - 9s), medium (10 - 29s) and long (≥ 30s) feeding bouts. The feeding bout ended when the snout of the piglet was out of the trough for a minimum of 5s. Exploratory behaviour towards the feed trough such as chewing the trough was not scored as eating. Daily/weekly eating activity per piglet was (semi-) quantified by summing the (minimum) number of seconds spent eating (where short, long and medium bouts counted as 5, 10 and 30 seconds respectively) from 2 days of age to weaning (at 28 days of age). All the observers were trained and instructed with the evaluating criteria to obtain homogeneous and accurate quantification of eating behaviour. Some parts of the videos were double checked by more than one person to evaluate consistency and observer dependent variations. This is to note that the eating behaviour measurements may have some degree of subjectivity and were taken as an “estimate” for the amount of eating per piglet and used as an indicative quantification of eating.

### Intestinal microbiota sampling and microbiota metataxonomic analysis

To investigate intestinal microbiota colonisation patterns, rectal swab samples were collected at eight time-points from 10 piglets per treatment group, following them till 2 weeks post-weaning. DNA extraction from rectal swab samples was performed by the repeated bead beating method (Yu and Morrison, 2004), followed by amplification of V3-V4 region of the bacterial 16S rRNA gene and Illumina (MiSeq) sequencing. Subsequently, the Illumina reads were imported and processed using the CLC Genomics Workbench version 11.01 (CLC bio, Arhus, Denmark). Detailed information about the sampling, DNA extraction, library preparation, sequencing and microbiota metataxonomic analysis can be found in **Supplementary file 1**.

### Performance data analysis

Piglets were weighed individually in all microbiota-sample-collection time-points pre- and post-weaning. Relative body weights of piglets (normalised to their weaning weights at day28) were calculated in the early post-weaning period to evaluate their body weight development during the weaning transition. To assess differences between the groups, Mann Whitney U-test and coefficient of variation (CV%) were calculated in GraphPad Software 8.1.1.

## RESULTS

### Dynamics of gut microbiome development in early-life of piglets

In this study, we first investigated the dynamics of intestinal microbiota development by analysing rectal swab samples collected from two days after birth (day2) to two weeks post weaning (day+14). All rectal swab samples were used to assess the microbiota development over time, irrespective of their allocation to the intervention group receiving pre-weaning fibrous feed (EF), or not receiving any additional feed (CON). 16S rRNA Illumina sequencing produced 5,013,336 reads after quality filtering, having a mean depth of 27,009 ± 5019 reads per sample.

Irrespective of the treatment, the porcine gut microbiota showed typical microbiota colonisation patterns with Firmicutes being the predominant phyla (60.01%), followed by Bacteroidetes (23.5%), Proteobacteria (8.04%), Actinobacteria (2.77%) and Fusobacteria (1.56%), together capturing 95.9% of total microbiome. Firmicutes and Bacteroidetes together accounted for 64-97% of total sequences across all ages. While the piglets aged from day2 to day+14 post-weaning, the relative abundance of Firmicutes increased (44% to 67%; *P* < 0.0001) and Bacteroidetes fluctuated pre-weaning eventually increasing post-weaning (19.9% to 29.2%; *P* < 0.0001) (**Supplementary figure 1**). The relative abundance of Fusobacteria significantly dropped from around 10% at day2 to less than 1% after 15 days of age (*P* < 0.05) almost disappearing over time (*P* < 0.0001). During the same time-period, the microbial (alpha) diversity strongly increased, especially during the pre-weaning period (**Figure 1A, Supplementary figure 2A)** and reached an apparent plateau post weaning. This is reflected by both the Chao 1 index (richness) that increased significantly from 2 days after birth (day2) up to day 5 post-weaning (day+5) as well as the Shannon index (richness and evenness) that increased gradually during the weaning period (day28) and remained stable afterwards.

**Figure 1:**
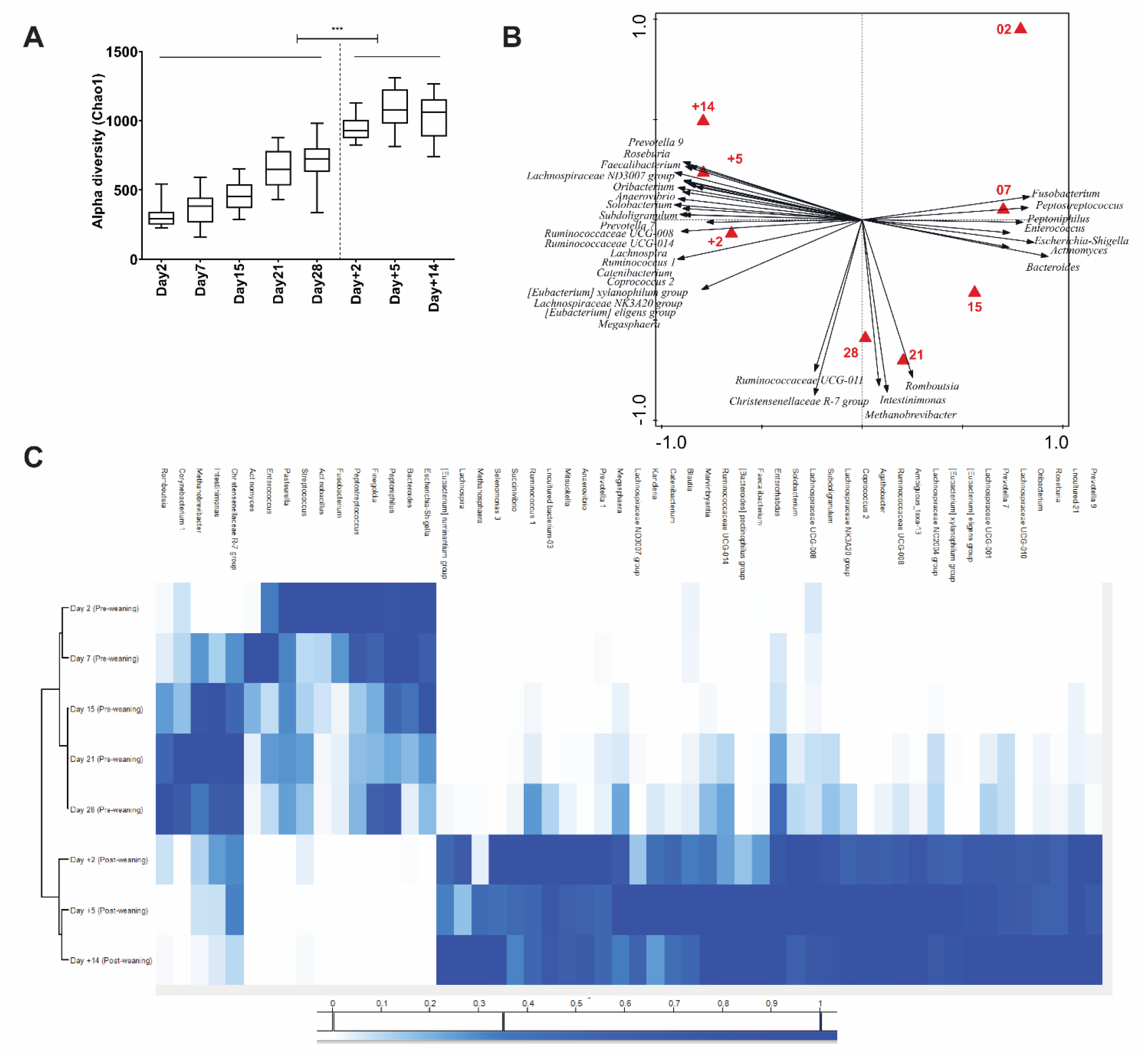
Age-associated intestinal microbiota dynamics pre- and post-weaning. **(A)** Alpha diversity (Chao1 bias corrected) display shifting diversity over time. **(B)** Redundancy analysis of age at genus level (explained variation = 28.22%, *P* = 0.002) with associated microbial groups at different ages. The age-related microbial groups visualized have minimum 35% fit on horizontal axis with a response score ≥ 0.7 in the biplot (obtained by projecting the taxa points perpendicular to the axes). **(C)** Heat map showing normalised relative abundance of the discriminative bacterial genera identified in redundancy analysis of age. Significant differences between time-points were assessed by student t test or Mann-Whitney U test (***: *P* < 0.001).

The impact of variables in the experimental set-up such as age, pen, gender and weaning, on the microbiota composition were investigated by redundancy (RDA) and partial redundancy (pRDA) analyses. The pen effect on microbiota was assessed by partial RDA (pRDA) corrected for other variables (age, treatment, gender) and revealed the expected (Frese et al., 2015) small but significant impact of pre-weaning pens on the intestinal microbiota (explained variation = 4.32%, *P* = 0.002). The effect of gender was evaluated by pRDA analyses for each time-point separately and did not reveal any gender-related influence on microbiota composition in the present experiment, which is analogous to what has been previously reported (Mach et al., 2015; Ke et al., 2019). A strong age-related effect (corrected for variables pen, gender, and treatment) on microbiome composition at genus level was observed (**Figure 1B**; explained variation = 28.22%, *P* = 0.002). The microbiota composition development as a function of age from day2 to day+14 displayed a horse-shoe shaped progression curve over time (**Figure 1B;** biplot with red triangles indicating different ages). Moreover, it allowed the identification of dominant microbial genera (black arrows) associated with certain time-points or parts of the overall timespan of the experiments. These microbial genera provided the markers for the time-dependent colonisation pattern in these piglets, irrespective of their treatment (EF or CON). For example, during the pre-weaning period, microbial groups such as *Actinomyces, Fusobacterium, Enterococcus, Peptostreptococcus, Bacteroides, Escherichia-Shigella* were abundant during the first few weeks of age as shown in heatmap representation (**Figure 1C**). On the other hand, genera like *Romboutsia, Intestimonas, Methanobrevibacter, Christensellaceae group R-7* emerged during the third or fourth weeks of age, but strikingly disappeared again post-weaning (**Figure 1C**). As expected, a sharp distinction was observed between the pre- and post-weaning microbiota, with several bacterial groups like *Prevotella, Faecalibacterium, Lachnospira, Roseburia, Subdoligranulum, Catenibacterium, Succinivibrio, Megasphaera, Coprococcus* particularly associated with post-weaning microbiota composition (**Figure 1B, 1C, Supplementary figure 2B**). Hierarchical clustering of age-related microbial genera (detected by RDA analyses) clearly separated the pre- and post-weaning associated microbiota, which is most probably governed by the sudden dietary shift, from pre-weaning mother’s milk to post-weaning solid (plant-based) feed. We observed a homogenising effect on the microbiota during the post-weaning period, which was characterised by comparable alpha diversity scores **(Figure 1A)**, closely clustered post-weaning time-points (**Figure 1B)** and lower Bray Curtis (PCoA) distance between post-weaning samples (**Supplementary figure 2C**).

### Impact of early-life (fibrous) feed intervention on microbiome development

#### Video observation shows eating behaviour increasing with time

Four pens (EF group) were provided with fibrous feed from two days of age in addition to suckling mother’s milk. We observed their eating behaviour around the feed trough using video recordings (0700 to 1900 hrs daily; eating time measured in seconds) for four weeks pre-weaning. The quantified eating behaviour scores for the EF piglets gradually increased as the piglets became older, reaching a clear amplified level of eating in the last week before weaning (**Figure 2A**). Eating behaviour scores during the first week of life were almost negligible, with piglets spending no- or only a very limited time exploring and eating the feed. However, during the second week of life, all the piglets started exploring and consuming the provided feed, and this behaviour increased during the third and increased strongly in the fourth week just before weaning. This individual eating behaviour was also supported by the previously reported (Middelkoop et al.) increase in feed intake at pen-level during the 4 weeks pre-weaning. However, despite this universal and prominent increase in eating behaviour over time, the quantified estimates of eating of individual piglets displayed considerable variation, analogous to the variation of feed-intake measurements per pen (Middelkoop et al.).

**Figure 2:**
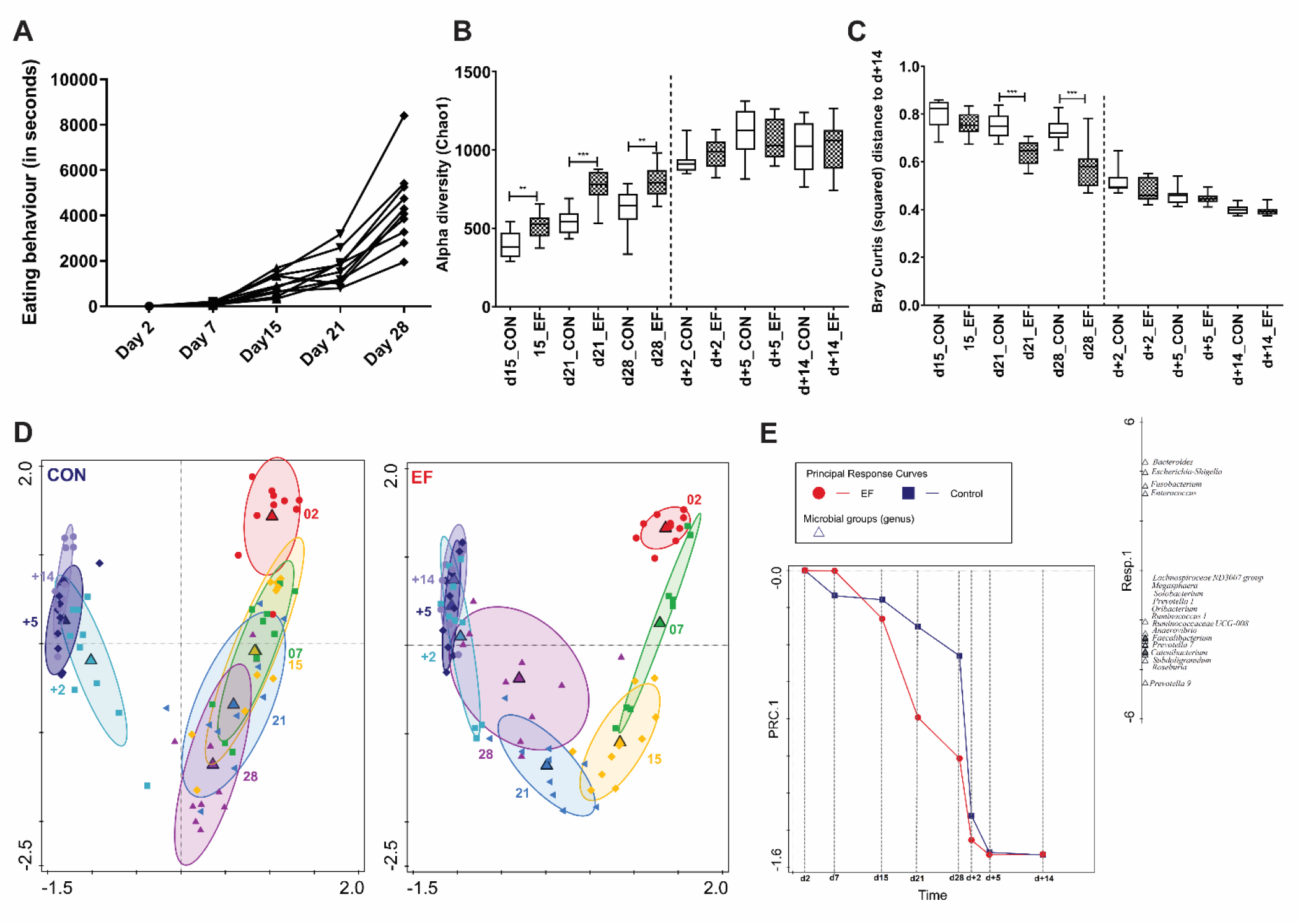
Microbiota colonisation development in early-fed (EF) and control (CON) group. **(A)** Eating behaviour (total eating seconds per week; n = 10) in EF litters over time pre-weaning. **(B)** Alpha diversity (Chao1 bias corrected) display diversity differences between the two groups from 15 days onwards. **(C)** Comparison of squared Bray Curtis index (distance between different samples with their d+14 time-point in individual piglets) between two groups. **(D)** Principal component analysis of microbiota in all ages (PC1 = 42.4%, PC2 = 12.3%), shown for the CON and EF group separately. **(E)** Principal response curve (PRC) analysis presenting changes in microbiota composition (genus level) across time for EF (red) and CON (green) group. The horizontal axis represents time and the vertical axis denotes PRC score values. Microbial response scores are shown on the right in a 1-dimensional plot. The combination of PRC score values and microbial response scores offers a quantitative interpretation and direction of change at different time-points. The effect of treatment (including its interaction with time) was significant according to Monte Carlo permutation test (*P* = 0.002). Significant differences between groups were assessed by student t test or Mann-Whitney U test (**: *P* < 0.01; ***: *P* < 0.001).

#### Early-life feeding accelerates microbiome colonisation maturation

One of the objectives of our study was to evaluate the impact of early-life feeding of fibrous feed on the microbiota composition, compared to the control group of piglets that did not receive any pre-weaning feed. Therefore, we investigated this effect using the microbiota composition information from 15 days of age onwards, because at the timepoints prior to day15, the piglets spent very little time eating solid feed (see above). Remarkably, the period with significant observed eating behaviour (day15 and onward) coincided consistently with higher levels of richness of the microbiota determined in the EF group relative to the CON group (**Figure 2B)**.

Moreover, in EF piglets, squared Bray Curtis distance (**Figure 2C**) of individual samples (day21 and day28) was significantly lower from their corresponding day+14, indicating proximity to a more “matured” state. The closeness of the EF group to the post-weaning microbiome, was also reflected in the increased ‘microbiome age’ at pre-weaning time-points (**Supplementary figure 3A**). However, this difference was no longer observed post-weaning, indicating that the microbiota diversity in the CON group rapidly caught-up with that of the EF piglets. These differences between EF and CON groups could be recapitulated in group-specific (EF or CON) PCA analysis (**Figure 2D**), which clearly demonstrated a considerable overlap of pre-weaning time-points day28, and to a smaller extent day21, with the post-weaning timepoints in EF piglets, whereas such overlap was completely absent in the CON piglets (**Figure 2D**). The EF treatment response over time was further investigated by principal response curve (PRC) analysis revealing a significant interaction of early feeding with time on the first constrained axis (*P* = 0.002; **Figure 2E**). In comparison to the CON group, the microbiota composition of EF group was enriched for several microbial genera (**Figure 2E**), including *Prevotella 9, Roseburia, Subdoligranulum, Catenibacterium, Faecalibacterium*, which are among the genera that are also typically among the microbial groups associated with the post-weaning microbiota in comparison to the pre-weaning microbiota (**Figure 1C, Supplementary figure 2B**). Obviously, day28 and day21 emerged as interesting time-points to be explored further to evaluate accelerated microbiome colonisation in the EF group, which is in agreement with the increased eating scores observed during these later pre-weaning time-points. The strongest discrimination between the EF and CON group was detected by RDA analysis at day28 (*P* = 0.002, **Figure 3A**), followed by day21 (*P* = 0.002; **Supplementary figure 3B**) and day15 (*P* = 0.04; **Supplementary figure 3C**). The microbial groups identified as main discriminants between the EF and CON group on day28 included those identified by PRC analysis, e.g., *Prevotella 9, Roseburia, Faecalibacterium, Megasphaera, Succinivibrio, Subdoligranulum, Catenibacterium, Coprococcus*, corroborating the enrichment of the post-weaning associated microbial genera in the EF group pre-weaning relative to the CON group. The maturation is also illustrated in hierarchical clustering of age-related microbial genera (detected by RDA analyses), where the EF group at day28 is clearly separated from the rest of the pre-weaning clusters (**Supplementary figure 4**). These microbial groups start colonising EF piglets from 21 and/or 28 days of age (**Figure 3C, Supplementary figure 5**). For example, *Prevotella 9*, the dominant genus of the post-weaning phase, started appearing from day21 in the EF group and continued to have significantly higher abundance at day28 (**Figure 3B**), compared to the CON group. RDA analyses at earlier time-points like day15 and day21 (**Supplementary figure 3B, C**) demonstrate the emergence of some microbes like *Succinivibrio, Ruminoccus 2* (day15 onwards) and *Prevotella 9, Megasphaera, Lachnospira, Subdoligranulum, Coprococcus* (day21 onwards) which persisted post-weaning.

**Figure 3:**
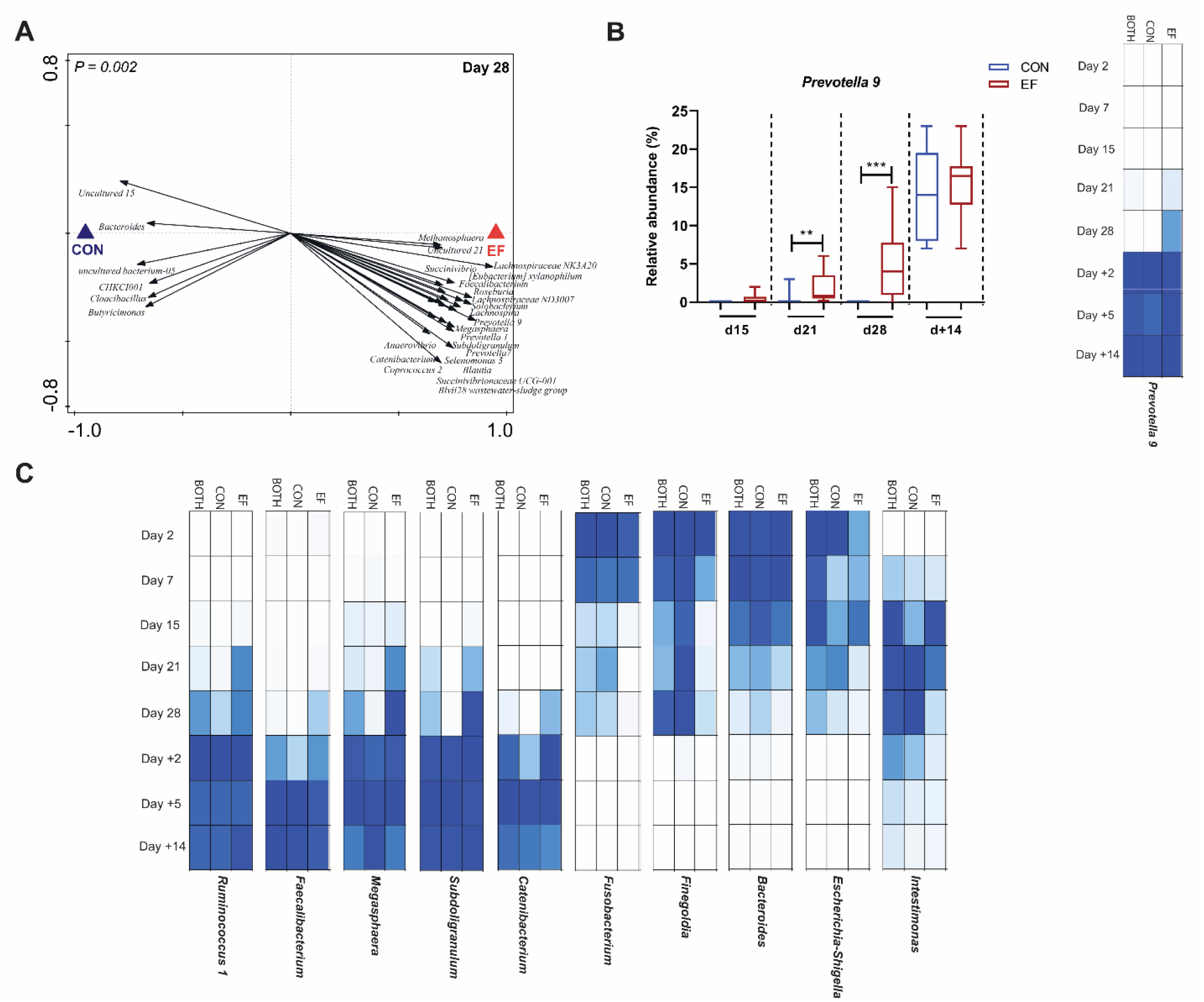
Accelerated microbiota maturation in early-fed (EF) group illustrated by emergence of post-weaning associated microbial groups in pre-weaning period. **(A)** Redundancy analysis of treatment at day28 (PC1 = 24.1%, PC2 = 19.7%; *P* = 0.002) with associated microbial groups in EF and CON groups (response score ≥ 0.65 on horizontal axis). **(B)** Changes in relative abundance of *Prevotella 9 in* EF and CON groups at pre- and post-weaning time-points. **(C)** Heat map of normalised relative abundance for representative individual genera at different ages showing appearance of post-weaning associated microbial groups and rapid loss of pre-weaning associated microbes, simultaneously. Significant differences between groups were assessed by student t test or Mann-Whitney U test (**: *P* < 0.01; ***: *P* < 0.001).

Furthermore, the EF group also displayed an accelerated loss of pre-weaning (day2-day7-day15-day21-day28)-associated microbial genera, like *Fusobacterium, Finegoldia, Intestimonas, Bacteroides* (**Figure 3C**), that is mostly found in the CON group (data not shown). Taken together, these results show that early feeding of fibrous diet accelerates the porcine microbiota colonisation, characterised by the earlier appearance and subsequent expansion of post-weaning associated microbial groups, in parallel with the more rapid disappearance of pre-weaning associated microbes. The pre-weaning microbiota differences induced in the EF group appeared to persist somewhat until five days post-weaning (**Supplementary figure 6A, B**), but disappeared and became undetectable two weeks post-weaning (day+14).

#### Quantitative relation between eating behaviour and microbiota

Since the accelerated microbiota colonisation patterns in the EF group coincided with the increased eating behaviour of individual piglets after two weeks of age (> 15 days), we investigated the quantitative relation between observed eating behaviour and the microbiota composition within the EF group.

Spearman correlation analysis was performed between eating behaviour (summed score per time-point) and squared Bray Curtis distance of individual piglets between a certain time-point and their day+14 “matured” time-point. It revealed a strong and significant negative correlation between these parameters, for both short term (last day; **Supplementary figure 7A**, last two days; r = - 0.798, *P* < 0.0001; **Figure 4**) and longer term (last week; **Supplementary figure 7B)** eating scores prior to the microbiota analysis timepoint. This indicates the consistency of eating behaviour in individual piglets, which is reflected in its association with both short-term and long-term scores. Since eating behaviour increased with age, the correlation analysis was done using different eating scores (ranging from last day – last week prior to the microbiota analysis timepoint). We found that inclusion of only ‘recent eating scores’ already quantitatively correlated with the magnitude of microbiota adaptation. Similarly, there was a strong positive association between microbiome age and eating behaviour (last two days; **Supplementary figure 7C**), fortifying the quantitative relation of eating behaviour and maturing microbiota.

**Figure 4:**
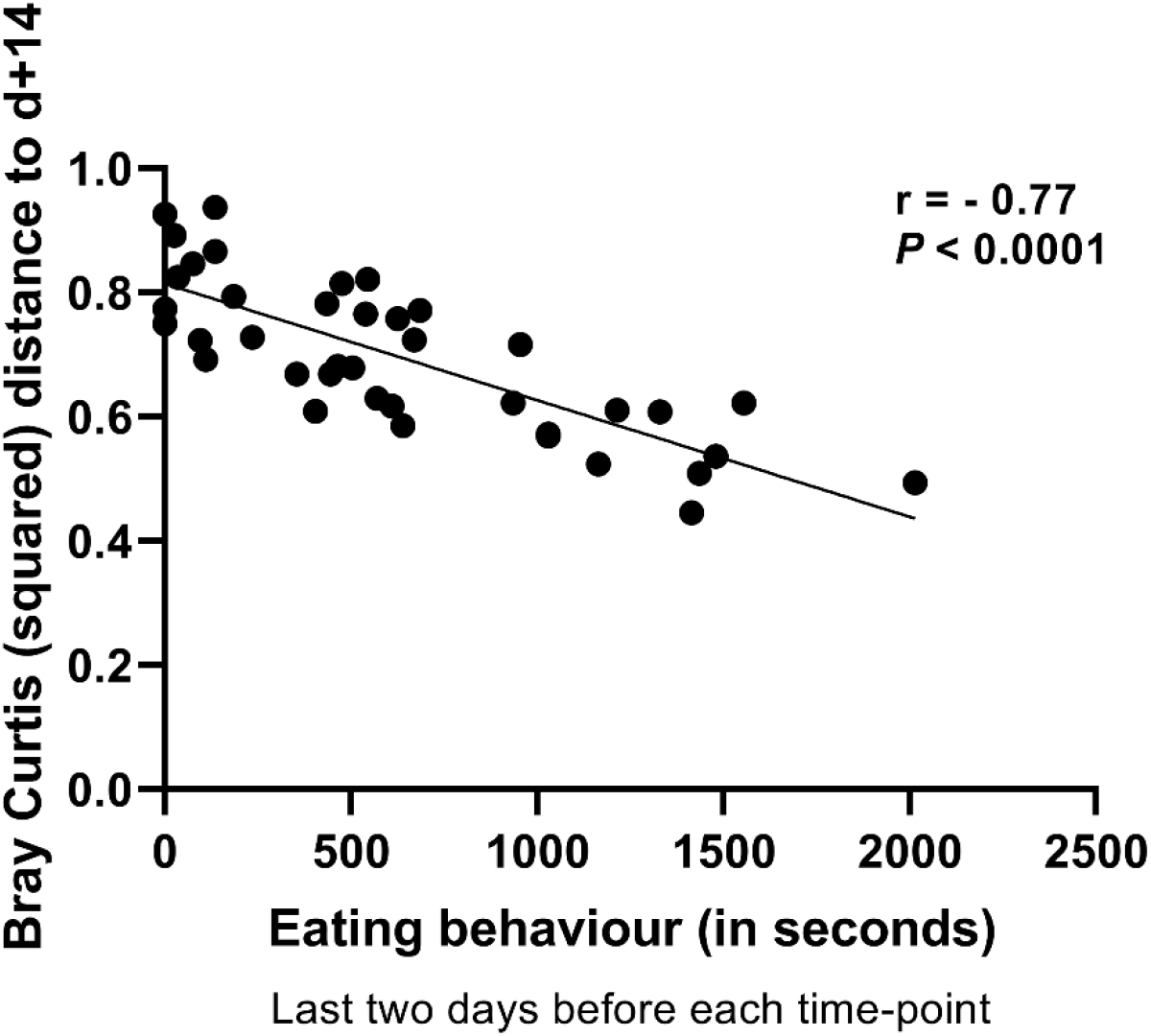
Spearman correlation between eating scores of individual piglets and squared Bray Curtis distance to their d+14 “matured” time-point. The last two days eating behaviour before each corresponding time-point was employed (r = - 0.77, *P* < 0.0001).

### Impact of early feeding on post-weaning body weight

To assess the impact of early feeding on post weaning performance, body weight of piglets was measured at three time-points day+2, day+5 and day+14 post weaning. The EF piglets tend to have a higher relative body weight compared to CON piglets at day+5 post weaning (**Figure 5**). Notably, we also observed a “smoother” transition for EF piglets in terms of relative body weight gain within the first few days post weaning, which was further supported by lower coefficients of variation and a consistent lack of body weight loss in piglets belonging to the EF group.

**Figure 5:**
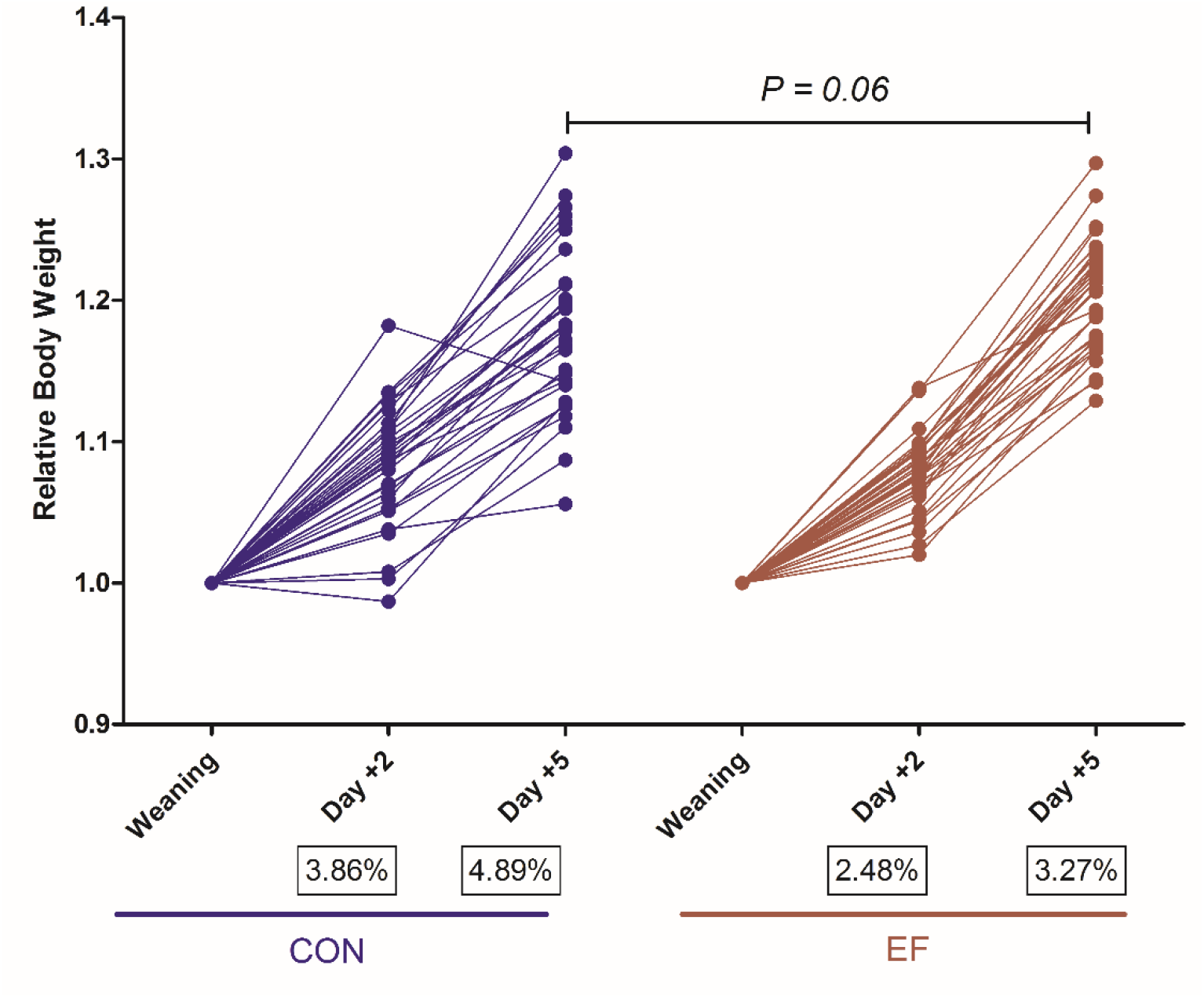
Comparison of relative body weight between the groups (n=32 per group) during the weaning transition (till 5 days post-weaning). Statistical comparisons between the groups were assessed by student t test or Mann-Whitney U test in GraphPad Software 8.1.1. The coefficient of variation (CV%) is indicated in black boxes below the labels.

## DISCUSSION

It is known that early-life perturbations can impact the intestinal microbiota, however only a few studies have utilised this “window of opportunity” to improve the weaning transition. In this study, we have assessed the age-associated microbiota development in young piglets and evaluated the impact of early-life feeding (pre-weaning access to a fibrous feed) on the gut microbiota colonisation and maturation over time (until 2 weeks post-weaning). Our results show that exposure to a fibrous diet in early-life accelerates the intestinal microbiota maturation pre-weaning, which is characterised by an increased microbial diversity, the earlier emergence and expansion of post-weaning-associated microbial groups and an enhanced loss of pre-weaning associated microbes. Importantly, analysis of individual piglets allowed us to establish that the magnitude of these microbiota compositional changes corresponds strongly with the individualized quantified eating behaviour during the pre-weaning period.

We observed a dynamic, age-related microbiota development with varied microbial groups associated at different time-points, and a discrete pre- and post-weaning associated microbiota. This is in line with multiple studies (Alain et al., 2014; Frese et al., 2015; Mach et al., 2015; Niu et al., 2015; Slifierz et al., 2015; Zhao et al., 2015; Guevarra et al., 2018; Li et al., 2018; Ke et al., 2019; Wang et al., 2019) which assessed bacterial communities in early-life of piglets, showing age as well as weaning as the driving factors in influencing microbiota development. The sudden shift in microbial communities after weaning is likely due to the abrupt dietary change from a highly palatable milk diet to a not-as-easily digestible plant-based solid feed. Firmicutes followed by Bacteroidetes were found to be the dominant phyla across all experimental time-points, and were particularly abundant post-weaning, where they accounted for more than 90% of all microbes detected. Although this is consistent with the majority of earlier studies (Kim et al., 2011; Alain et al., 2014; Mach et al., 2015; Niu et al., 2015; Chen et al., 2017; Holman et al., 2017; Ke et al., 2019; Wang et al., 2019), there are a few reports with Bacteroidetes (Lu et al., 2014; Guevarra et al., 2018) or Proteobacteria (Slifierz et al., 2015; Zhao et al., 2015) as the pre-dominant phyla. Analogously, the decreasing abundance of Fusobacteria observed during the first weeks of life has also been reported by several other studies (Alain et al., 2014; Niu et al., 2015; Slifierz et al., 2015; Ke et al., 2019), although there are also studies where this bacterial group was not detected at all (Frese et al., 2015; Guevarra et al., 2018). Such differences in taxonomic identification could, to some extent, be due to the different primers and variable regions of the 16S rRNA gene that were targeted in these metataxonomic investigations, which has been shown to capture different community structures (Albertsen et al., 2015; Holman et al., 2017), disallowing an accurate comparison between different studies. It should be noted that the other phyla we detected in this study, like Actinobacteria, Spirochaetes, Synergistetes, were consistently present at low relative abundance levels, which may contribute to their large fluctuations among studies. Despite these variations among studies, and the various confounding factors among study designs and conditions, including pig genetics, environmental conditions, piglets age at weaning, sampling procedures, sample processing and sequence analysis methods, there is a striking congruency in the biological interpretation of differences between pre-and post-weaning associated microbes in most studies (Alain et al., 2014; Frese et al., 2015; Mach et al., 2015; Slifierz et al., 2015). This is largely in agreement with our observation in the present study and pinpoints the early intestinal colonisers belonging to *Bacteroides, Escherichia-Shigella* and exemplifies *Prevotella* as a prominent microbe in the typical post-weaning microbiota together with species belonging to *Roseburia, Faecalibacterium, Blautia, Subdoligranulum*. The drastic increase in relative abundance of *Prevotella* post-weaning (from less than 3% to more than 18% of the total community) is likely due to the established capacity of the members of the genus to produce enzymes that can degrade the plant polysaccharides (Flint and Bayer, 2008; Ivarsson et al., 2014), which are prominent constituents of the plant-based weaner diet. Similarly, the current observation of the increasing microbial diversity (Chao and Shannon) with age, which eventually reached a plateau post-weaning (within 2 weeks), is in good accordance with several previous studies (Alain et al., 2014; Frese et al., 2015; Niu et al., 2015; Slifierz et al., 2015; Chen et al., 2017; Ke et al., 2019; Wang et al., 2019), although opposite diversity development patterns have been reported in a few studies as well (Guevarra et al., 2018; Li et al., 2018). This discrepancy might be due to variations in weaning age (e.g., the last two studies employed 21 days as weaning age) as well as the post-weaning sampling time-point (e.g., the last two studies had a shorter post-weaning sampling time-point; within a week). Microbiota metrics like alpha diversity, Bray Curtis distance, as well as hierarchical clustering clearly separated samples taken pre- and post-weaning, with the shorter-term post-weaning samples (2 and 5 days post-weaning) being remarkably close to the last post-weaning timepoint (14 days post-weaning). This suggests that the porcine microbiota rapidly evolves post-weaning, towards an apparently homogeneous and stable microbiome structure within 2 weeks post-weaning, reflecting the crucial role of diet in dictating the microbiome adaptation during the weaning transition. These diet driven effects clearly overrule various pre-weaning factors that contribute to the higher microbiome variation observed prior to weaning, such as litter-specific genetic diversity, distinct consequences of vertical transmission from the sow, and/or variations in sow milk composition. Taken together, the present study confirms and refines various earlier observations of the microbiome development during early-life stages in piglets and exemplifies the major impact of the weaning transition that leads to the establishment of a homogenous, rich and stable microbiota composition post-weaning.

The central aim of the present study was to investigate whether provision of pre-weaning fibrous diet would impact the microbiome development in suckling piglets. Quantitative eating scores of individual early-fed (EF) piglets rapidly increased over time, reaching the highest eating score in the last week prior to weaning. This is in agreement with creep feeding studies (Fraser et al., 1994; Bruininx et al., 2002) which provided solid feed preweaning and found that piglets consumed about 60-80% of total feed in the last week before weaning (4 weeks of age). Unlike traditional creep feeding diets, in this study, the pre-weaning diet was designed to contain a mixture of soluble and insoluble fibres, that can potentially stimulate the large intestinal microbiota. We could show that piglets which consumed this fibrous diet pre-weaning displayed an accelerated microbiota maturation compared to the control (CON) piglets suckling sow’s milk only. Acceleration of the microbiota development in the EF piglets was observed by a higher alpha diversity compared to CON piglets from 15 days of age until weaning, coinciding with the observation of increased eating scores from 15 days of age, thereby suggesting the relation between feed consumption and these changes. Moreover, the EF piglets’ microbiota resemblance to the post-weaning microbiota increased during the last two weeks of pre-weaning period, an effect that was not noticeable in the CON piglets. This effect in the EF piglets was most prominently reflected by the emergence of increasing relative abundances of typical post-weaning associated genera like *Prevotella, Roseburia, Faecalibacterium, Ruminococcus, Megasphaera, Catenibacterium and Subdoligranulum*. Additionally, these effects coincided with the accelerated disappearance of typical pre-weaning associated microbes such as *Fusobacterium, Finegoldia, Bacteroides, Eschechichia-Shigella*, which did occur at a much slower rate in the CON piglets. Previous studies have occasionally found similar microbial groups on consumption of solid feed or specific dietary fibres in suckling piglets (Berding et al., 2016; Zhang et al., 2016; Schokker et al., 2018; Wang et al., 2019). Importantly, the degree of resemblance observed was quantitatively correlated to the amount of feed consumed pre-weaning, strengthening the relation between feed-consumption and the acceleration in microbiota development.

Furthermore, in the present study, the EF piglets showed a tendency of higher relative body weight at 5 days after weaning, compared to CON piglets. Intriguingly, we observed a smoother transition post-weaning in EF piglets, with a lower variation (CV) and lack of negative trendlines in body weight development from weaning to day+2, thus reflecting consistency in post-weaning performance. Microbial fermentation of undigested fibres results in SCFA formation, creating a niche for bacterial groups that possess saccharolytic properties. For example, *Prevotella* that represents a group of strictly anaerobic bacteria, is reported as the dominant genus in the large intestine of pigs (Leser et al., 2002; Liu et al., 2012; Wang et al., 2019), especially after the introduction of fibrous solid feed post-weaning. The discriminating microbial groups that we observed in EF piglets are mostly fibre-degrading, anaerobic, SCFA producers. One of the expected differences between the two groups is that the EF piglets may have higher SCFA concentrations in distal part of the intestine, contributing to more SCFA exposure in intestinal mucosa, thereby stimulating intestinal development towards adaptation for better handling of post-weaning diet by properly digesting and acting on nutrients. This could potentially explain the smoother transition during the first five days post-weaning. Further, the CON group quickly ‘catches up’ post-weaning after exposure to weaner diet, towards a homogeneous microbiota observed in both groups at day+14 post-weaning. However, they made a ‘longer distance’ in that short time period, which is possibly reflected in their less smooth and higher variation in body weight development. In future studies, enlarging sample size could lead to detection of significant post-weaning weight development effect, which was observed as a trend in the current study.

In conclusion, this study confirms age-related microbiota dynamics in early-life stages (first weeks of life) and their progression over time in neonatal piglets. The present study illustrates the impact of early-life feeding of fibrous diet on gut microbiota, showing an increased microbiome maturation or higher “microbiome age” in early-fed piglets at pre-weaning stages. Further, we observed a strong quantitative association between eating behaviour and microbiota, suggesting that the piglets who spent more time at the feeding trough had higher abundance of ‘accelerated’ microbial groups. This indicates the importance of developing pre-weaning strategies (fibrous solid diet) to enhance eating behaviour in suckling piglets for increased eating and more matured microbiota. Overall, these findings emphasises the importance of early-life “window of opportunity” for modulation of the gut microbiota development and could aid in the development of optimal nutritional strategies to support the timely gut colonisation of relevant microbial taxa in the gradually diversifying neonatal gut.

## ACKNOWLEDGEMENTS

This study is part of research programme “Genetics, nutrition and health of agricultural animals” with project number 868.15.010, which is financed by the Netherlands Organization for Scientific Research, Cargill Animal Nutrition and Coppens Diervoeding. The authors want to thank the technicians and students of the Host-Microbe Interactomics group and Adaptation Physiology group (Wageningen University & Research) who helped during the animal experiment, sample processing and video observations. We would like to acknowledge the personnel and trainees of the animal research facilities in Wageningen (Carus) for taking care of the animals and for their technical assistance. The authors are grateful to Tamme Zandstra for producing the experimental feed and to FrieslandCampina DOMO (Amersfoort, The Netherlands) for provision of Vivinal GOS. We also want to thank Jos Boekhorst for his inputs in microbiota analysis employing Canoco software.

## Supplemental information

### Supplementary file 1

To investigate intestinal microbiota colonisation patterns, rectal swab samples were obtained from piglets by inserting a sterile cotton swab (Puritan Medical, Guilford, ME USA; Cat Number-25-3306-U) 20–30 mm into the rectum and rotating the swab against the bowel wall for a minute before placing it into a 5ml eppendorf tube. The samples were kept on ice during transport to the laboratory and stored at −20°C until further processing. The selection of piglets (n=10 per treatment group) were made by the following criteria: (a) no antibiotic treatment (b) no pre-weaning diarrhoea (c) close to average weight of the treatment group (d) equal male to female ratio. A total of 160 swab samples were collected, repeatedly from the same piglets at five time-points pre-weaning (2, 7, 15, 21, 28 days of age) and three time-points post-weaning (+2, +5 and +14 days of age). At 2 and 7 days of age, samples were collected fixed to their birth date, thereafter all samples were taken on the same day for all piglets fixed to the day of weaning. However, one sample (from EF group at 7days of age) was unsuccessful at the sequencing step and therefore could not be included in the analysis. The EF piglets selected for microbiota analysis belonged to four pens (or litters) before weaning (2-3 piglets selected per pen), and two pens had to be omitted from microbiota analysis due to uterine infection of the sow and subsequent antibiotic treatment during the suckling phase.

DNA extraction from rectal swabs was performed by the repeated bead beating method (Yu and Morrison, 2004) using QIAamp DNA Stool Mini Kit (Qiagen, Hilden, Germany) according to the manufacturer’s instructions. 500 μl of lysis buffer was added to the 5ml eppendorf tube (holding swab) to obtain swab solution, which was used as a starting material for DNA extraction. The quality and quantity of extracted DNA samples were checked by gel electrophoresis (only representative samples) and Nanodrop DeNovix DS-11 Spectrophotometer (DeNovix Inc., Wilmington, DE USA) respectively. The DNA template was used for amplifying the V3-V4 region of the bacterial 16S rRNA gene using V3F primer (5′-CCTACGGGNGGCWGCAG-3′) and V4R primer (5′-GACTACHVGGGTATCTAATCC-3′), 5′-extended with extension-PCR-adapters 5′-TCGTCGGCAGCGTCAGATGTGTATAAGAGACAG-3′ and 5′-GTCTCGTGGGCTCGGAGATGTGTATAAGAGACAG-3′ respectively. PCR Amplification (Bio-Rad C1000 thermal cycler, Bio-Rad Laboratories, Veenendaal, The Netherlands) of the 16S rRNA V3-V4 region was completed in a 50 μl reaction volume consisting of 5 μl 10× KOD buffer (Toyobo, Japan), 3 μl MgSO4 (25 mM) 5 μl dNTPs (2 mM each), 1.5 μl V3F primer [10 μM (Eurogentec, Luik, Belgium)], 1.5 μl V4R primer [10 μM, (Eurogentec, Luik, Belgium)], 1.0 μl (0.02 U/μl) KOD hot start DNA polymerase (Toyobo, Japan) and 10 ng (minimum) of template DNA. The amplification conditions included a single initiation cycle of 95°C for 2 min, followed by 25 amplification cycles encompassing denaturation at 95°C for 20 s, annealing at 55°C for 10 s, and elongation at 70°C for 15 s, and was completed by a single elongation step at 72°C for 5 min. Amplicons were purified using MSB Spin PCRapace (STRATEC Molecular, Germany) and were sequenced at BaseClear BV (Leiden, The Netherlands) using (paired-end) Illumina MiSeq system. Purified amplicons were subjected to extension-PCR using barcoded Illumina universal index sequencing adapters prior to sequencing. The Illumina MiSeq system generated FASTAQ sequence files using the bcl2fastq2 version 2.18 and these sequences were subjected to quality control based on Illumina Chastity filtering and FASTQC quality control tool version 0.11.5. Subsequently, a BaseClear in-house filtering protocol was applied for removal of reads containing adapters (up to minimum read length of 50bp) and/or PhiX control signal, to generate the FASTAQ data file used for microbiota analysis.

Illumina reads were imported into the CLC Genomics Workbench version 11.01 and were processed using the CLC Microbial Genomics Module version 2.5.1 (CLC bio, Arhus, Denmark). The paired end reads were merged into one high quality representative by CLC Workbench (Mismatch cost = 1, Minimum score = 40, Gap Cost = 4, Maximum unaligned end mismatches = 5). The CLC pipeline was used for primer and quality trimming (Trim using quality scores = 0.05; Trim ambiguous nucleotides: maximum number of ambiguities = 2; Discard reads below length = 5). The remaining high quality sequences were clustered into operational taxonomic unit (OTUs) at 97% identity threshold using SILVA database v132 (released on Dec 13, 2017) (Quast et al., 2013). OTUs lower than 2 reads (Minimum combined count = 2) were excluded from the analyses. To achieve even sequencing depth between samples, the OTU table was rarefied to 14,000 reads for calculation of alpha and beta diversity indices. Principal component analysis (PCA; unsupervised), redundancy analysis (RDA; supervised) and partial redundancy analysis (pRDA; supervised) were performed using CANOCO 5 (Microcomputer Power, Ithaca, NY, USA) according to manufacturer’s instructions (Braak and Smilauer, 2012). To evaluate the impact of environmental variables like age, pen, gender and treatment separately, RDA and pRDA analyses were performed, the latter analyses (pRDA) allowing us to correct for other covariates in the data as described in Canoco 5 manual (Braak and Smilauer, 2012). The pen-related impact was assessed separately per time-point and in pre- and post-weaning phases to avoid confounding age and weaning-related differences. The microbial groups detected by RDA analyses were selected by their response scores in the biplot (obtained by perpendicular projection of the taxa arrows on the X axis) and were further filtered by the condition of having ≥ 0.1% average relative abundance in at least one of the time-points (for biological relevance). The median relative abundance of selected microbial groups was calculated for each of the time-points in 20 piglets (assessing age) or 10 piglets (assessing treatment groups). Relative abundance of age-related microbial groups were visualised by heat maps by normalising by the highest median of all timepoints (scaling from 0 to 1), in Perseus software (Tyanova et al., 2016). Euclidean distance was utilized to measure the distance and clustering was conducted using the average linkage method. Time dependent treatment effect were assessed by using a linear ordination method known as principal response curves (PRCs) in Canoco 5, which is based on redundancy analysis (RDA), adjusted for overall changes in microbial community response over time (Van Den Brink and Ter Braak, 1999). The principal component is plotted against time, yielding a principal response curve of the community for each treatment. The PRC method also shows microbial response scores by a 1-dimensional plot, thereby enabling quantitative interpretation of treatment effect towards the microbial species level. Statistical significance was evaluated by Monte Carlo permutation procedure (MCPP) with 499 permutations. Response-variable based case scores (CaseR) was extracted from redundancy analysis of age, which was defined as their ‘microbiome age’ as they correspond to the individual sample position in the progressing-age-ordination plot. To evaluate the quantitative relationship between observed eating behaviour and magnitude of the microbiota change, a non-parametric spearman correlation was performed between eating scores (in seconds) and squared Bray Curtis distance of piglets or ‘microbiome age’ between individual time-points and their corresponding day+14 time-point in GraphPad Software 8.1.1 (California, USA, www.graphpad.com). Differential abundance analysis between the CON and EF group was implemented in CLC Genomics Workbench (edgeR test) from day 15 onwards, as shown in **Supplementary table 2**. Comparison of the specific taxa relative abundance and diversity indices between treatments were performed by Mann Whitney U-test whereas comparison among time-points were assessed by one-way ANOVA (Kruskal-Wallis statistic) using a Dunnett’s test for multiple comparison using GraphPad Software 8.1.1. The level of statistical significance was set at *P* < 0.05 with a trend defined as 0.1 > *P* ≥ *0*.*05*.

**Supplementary table 1:**
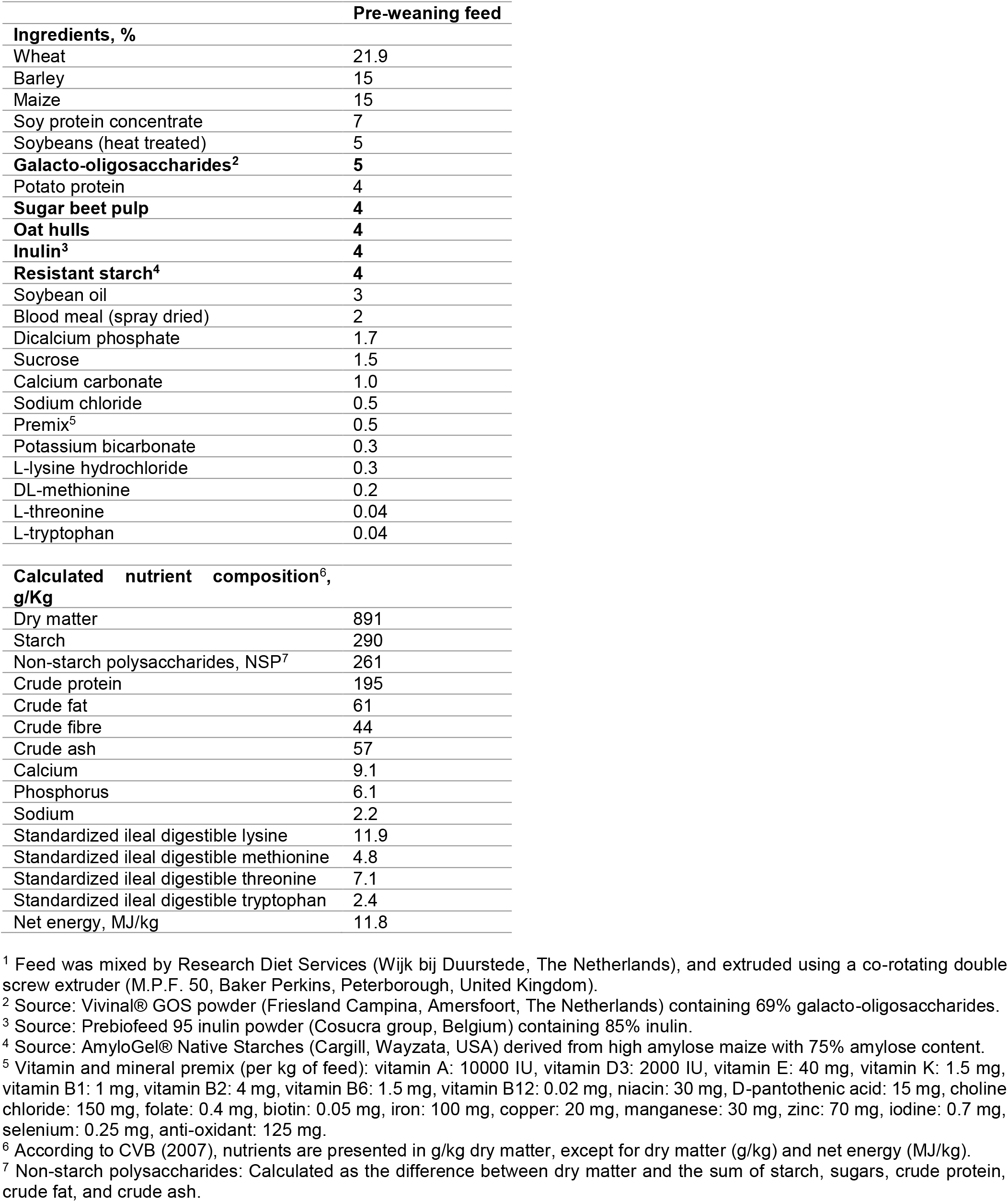
Ingredients and calculated nutrient composition of the pre-weaning fibrous feed^1^.

**Supplementary figure 1:**
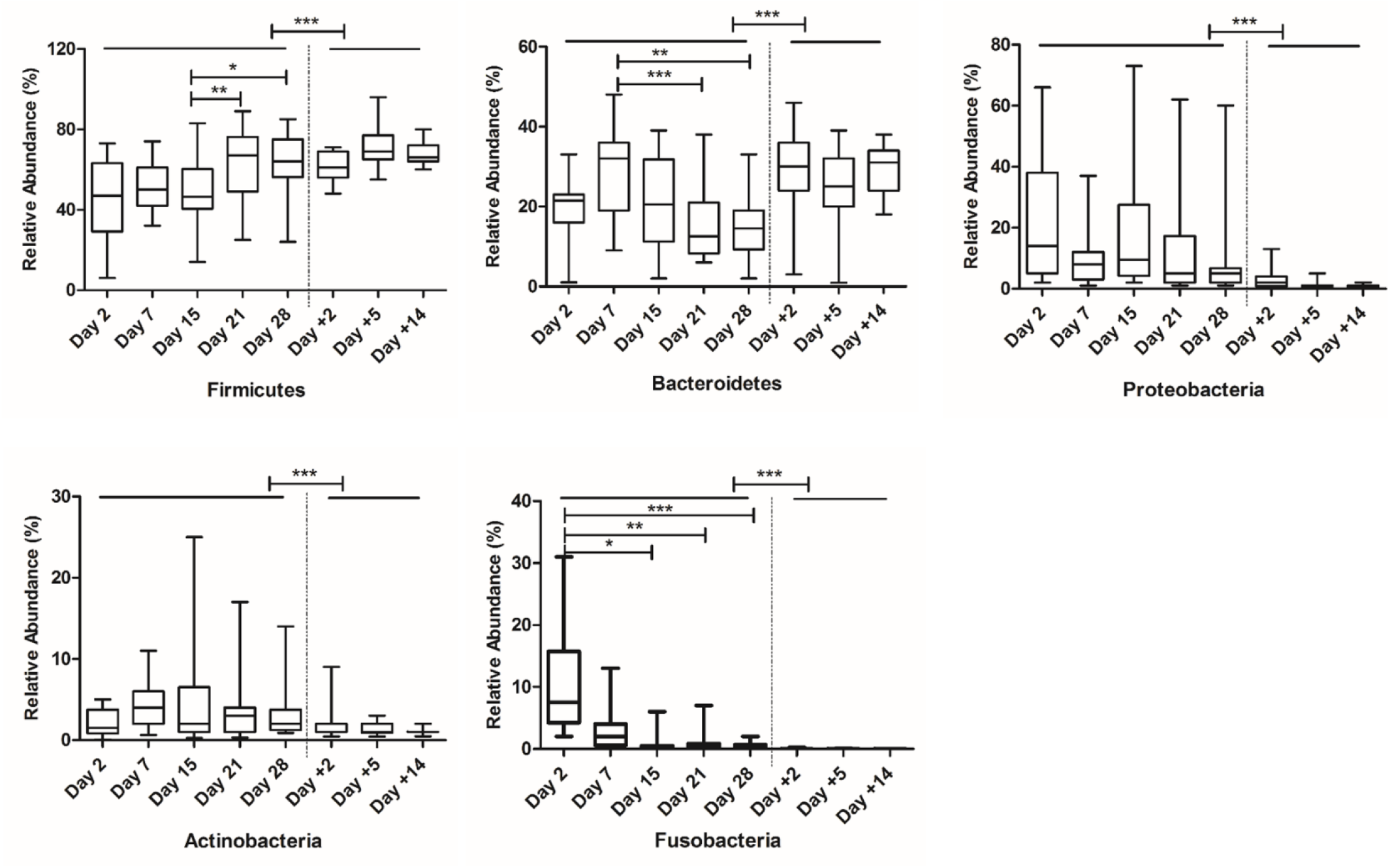
Age-related microbiota dynamics showing relative abundance of taxa (phyla level) at pre- and post-weaning time-points. Statistical differences were assessed by Mann Whitney U-test or one-way ANOVA (Kruskal-Wallis statistic) using a Dunnett’s test for multiple comparison. (*: *P* < 0.05; **: *P* < 0.01; ***: *P* < 0.001).

**Supplementary figure 2:**
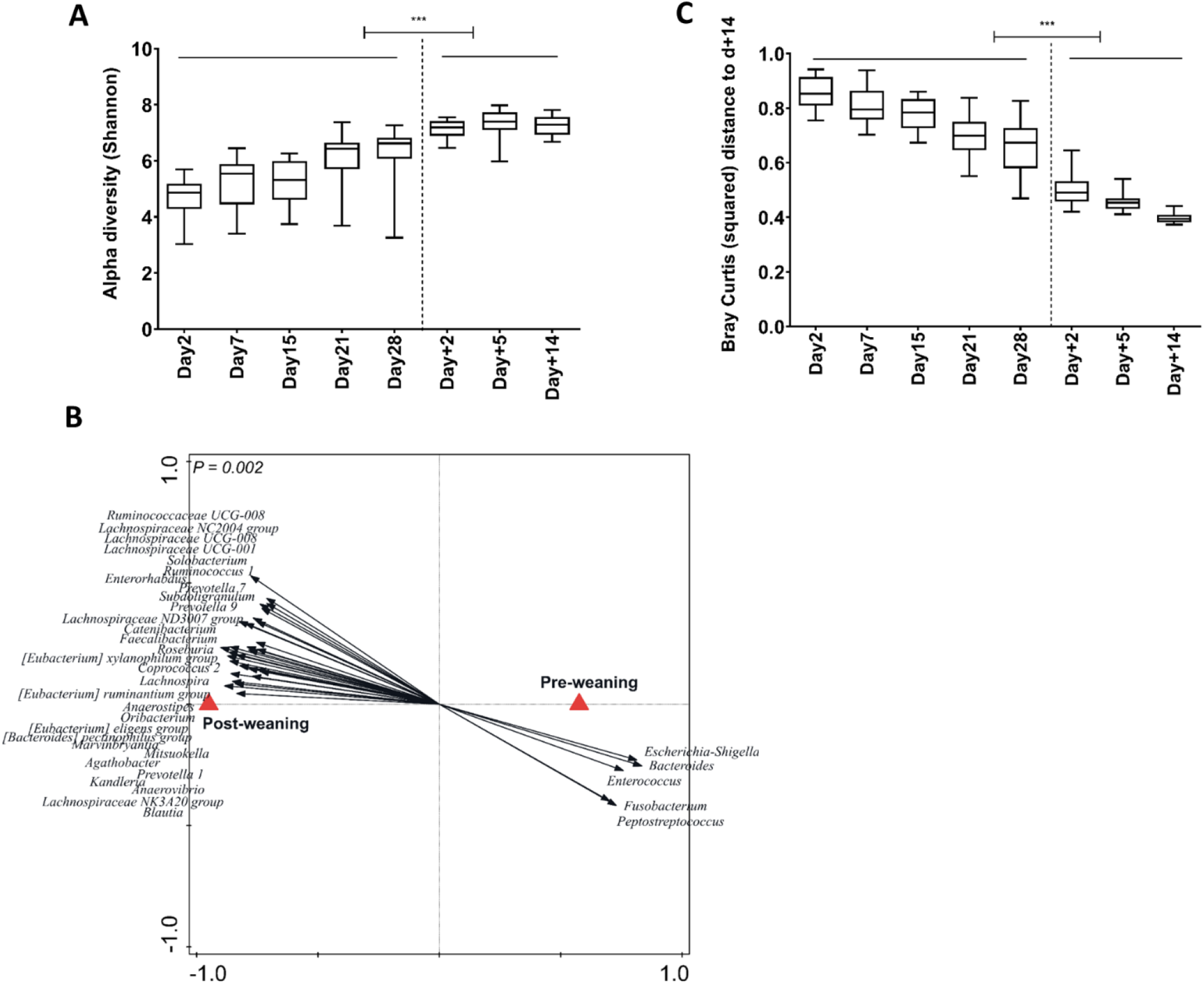
Age-driven intestinal microbiota progression in pre- and post-weaning period. **(A)** Alpha diversity (Shannon index) showing increasing diversity over time. **(B)** Redundancy analysis of weaning (explained variation = 32.4%; *P* = 0.002) showing both pre- and post-weaning associated microbiome. The microbial groups visualized have minimum 40% fit on horizontal axis with a response score ≥ 0.7 in the biplot (obtained by projecting the taxa points perpendicular to the horizontal axis). **(C)** Bray Curtis (squared) distance between pre- and post-weaning time-points. Significant differences between groups were assessed by student t test or Mann-Whitney U test (***: *P* < 0.001).

**Supplementary figure 3:**
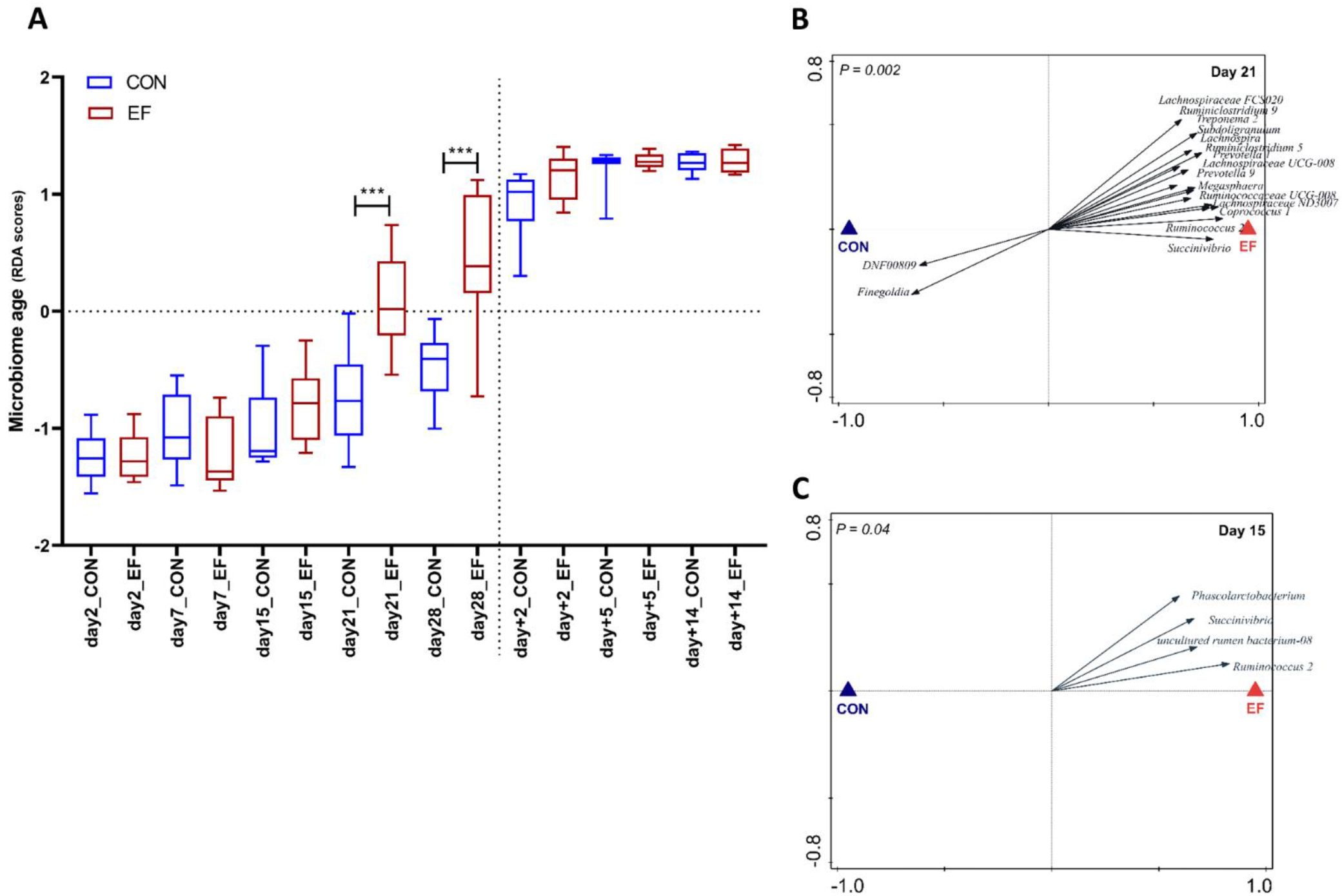
**(A)** Comparison of the developing ‘microbiome age’ between early-fed (EF) and control (CON) piglets. Response-variable based case scores (CaseR) that was extracted from redundancy analysis of age, was defined as ‘Microbiome age’ as they correspond to the individual sample position in the progressing-age-ordination plot. Significant differences between groups were assessed by student t test or Mann-Whitney U test (***: *P* < 0.001). Redundancy analysis of treatment at pre-weaning time-points: **(B)** day21 (PC1 = 15.6%, PC2 = 18.8%; *P* = 0.002) and **(C)** day15 (PC1 = 8.7%, PC2 = 24.3%; *P* = 0.04), with associated microbial groups in EF and CON groups (response score ≥ 0.6 on horizontal axis).

**Supplementary figure 4:**
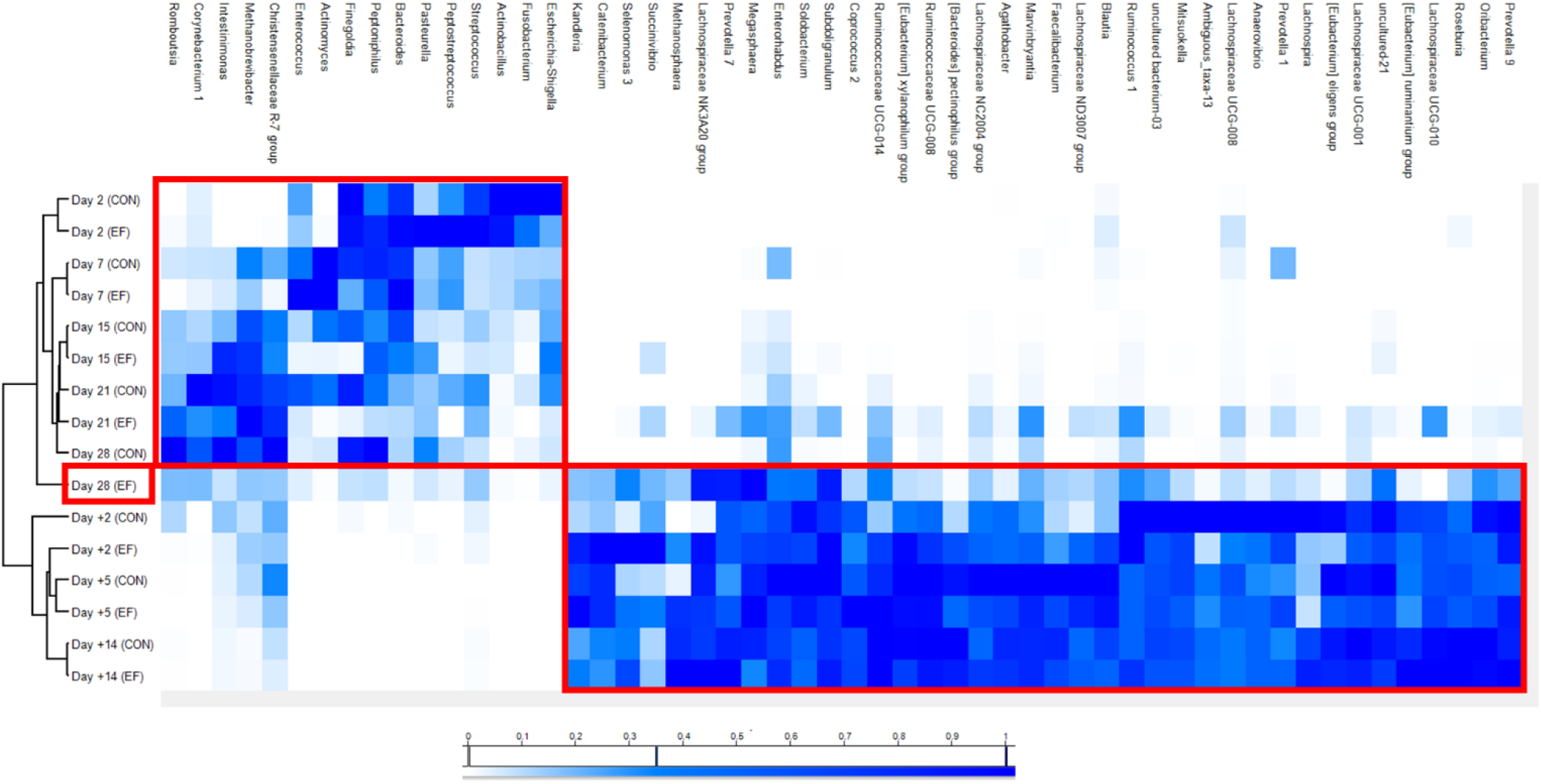
Hierarchical clustering of early-fed (EF) and control (CON) groups at different ages, using normalised relative abundance of the discriminative microbial genera (identified in redundancy analysis of age). The EF group at day28 clearly separates from the other pre-weaning groups and is closer to the post-weaning microbiome.

**Supplementary figure 5:**
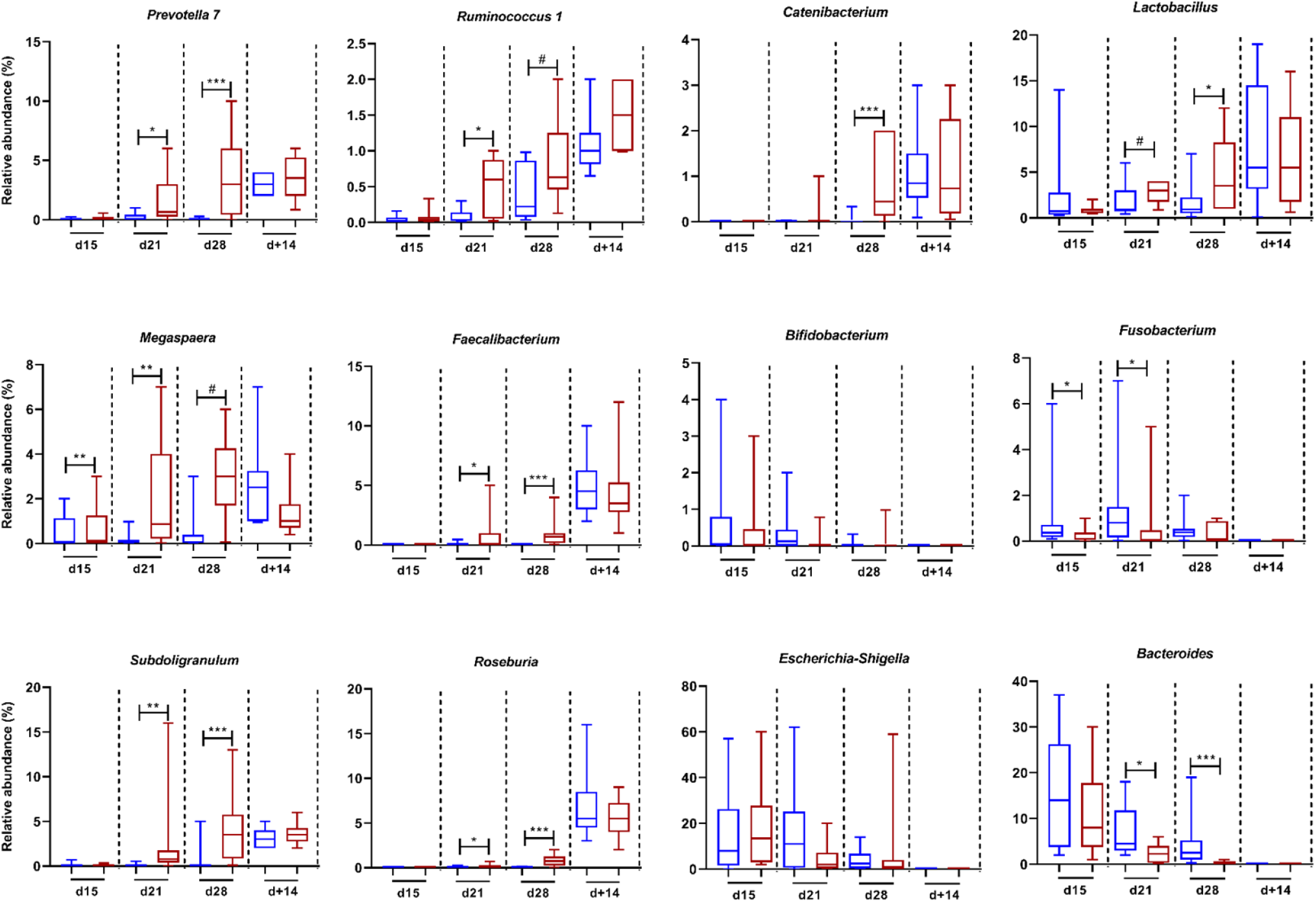
Changes in relative abundance of representative microbial groups (shown in box plots) in early-fed (EF; red) and control (CON; blue) groups at pre- and post-weaning time-points. Statistical differences were assessed by Mann Whitney U-test (non-parametric) or t test (#: 0.1 > *P* ≥ 0.05; *: *P* < 0.05; **: *P* < 0.01; ***: *P* < 0.001).

**Supplementary figure 6:**
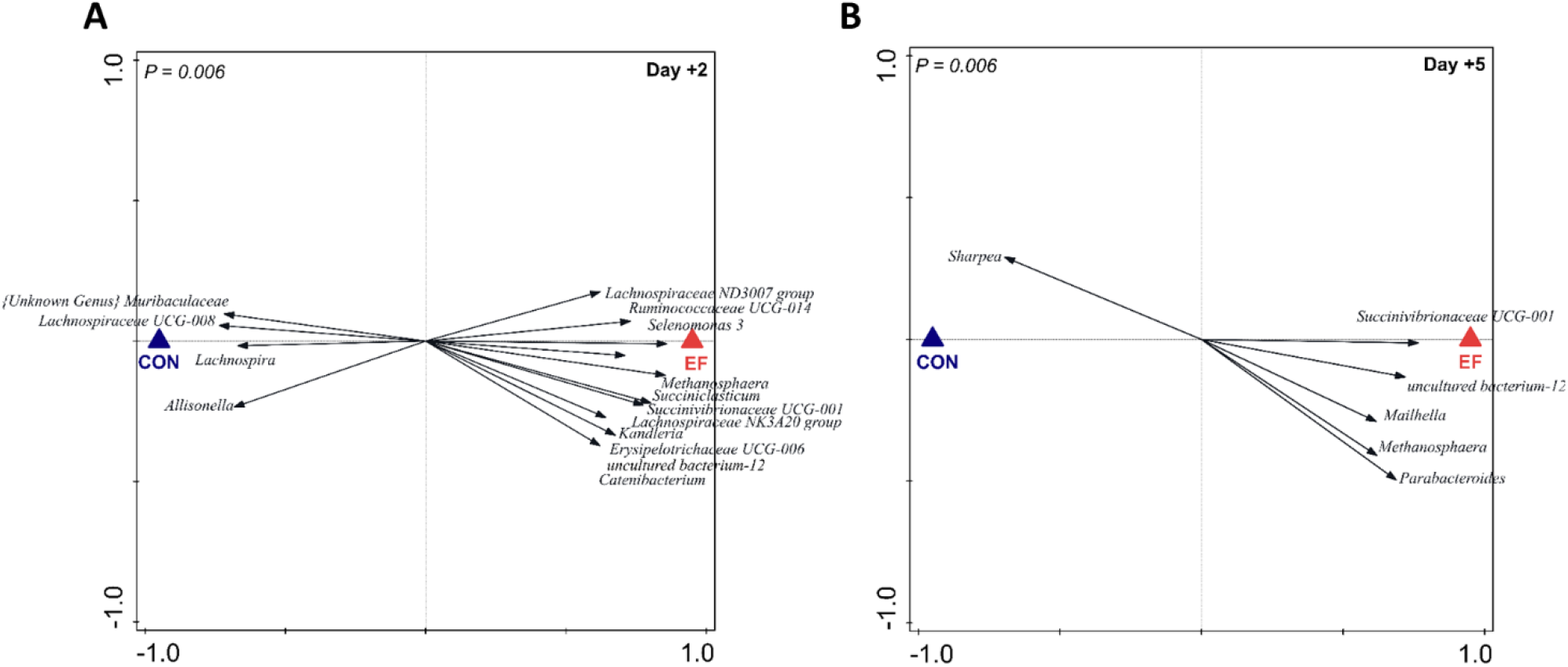
Redundancy analysis of treatment at post-weaning time-points **(A)** day+2 (PC1 = 16.01%, PC2 = 25.6%; *P* = 0.006) and **(B)** day+5 (PC1 = 10.8%, PC2 = 20.8%; *P* = 0.006) with associated microbial groups in early-fed (EF) and control (CON) groups (response score ≥ 0.6 on horizontal axis).

**Supplementary figure 7:**
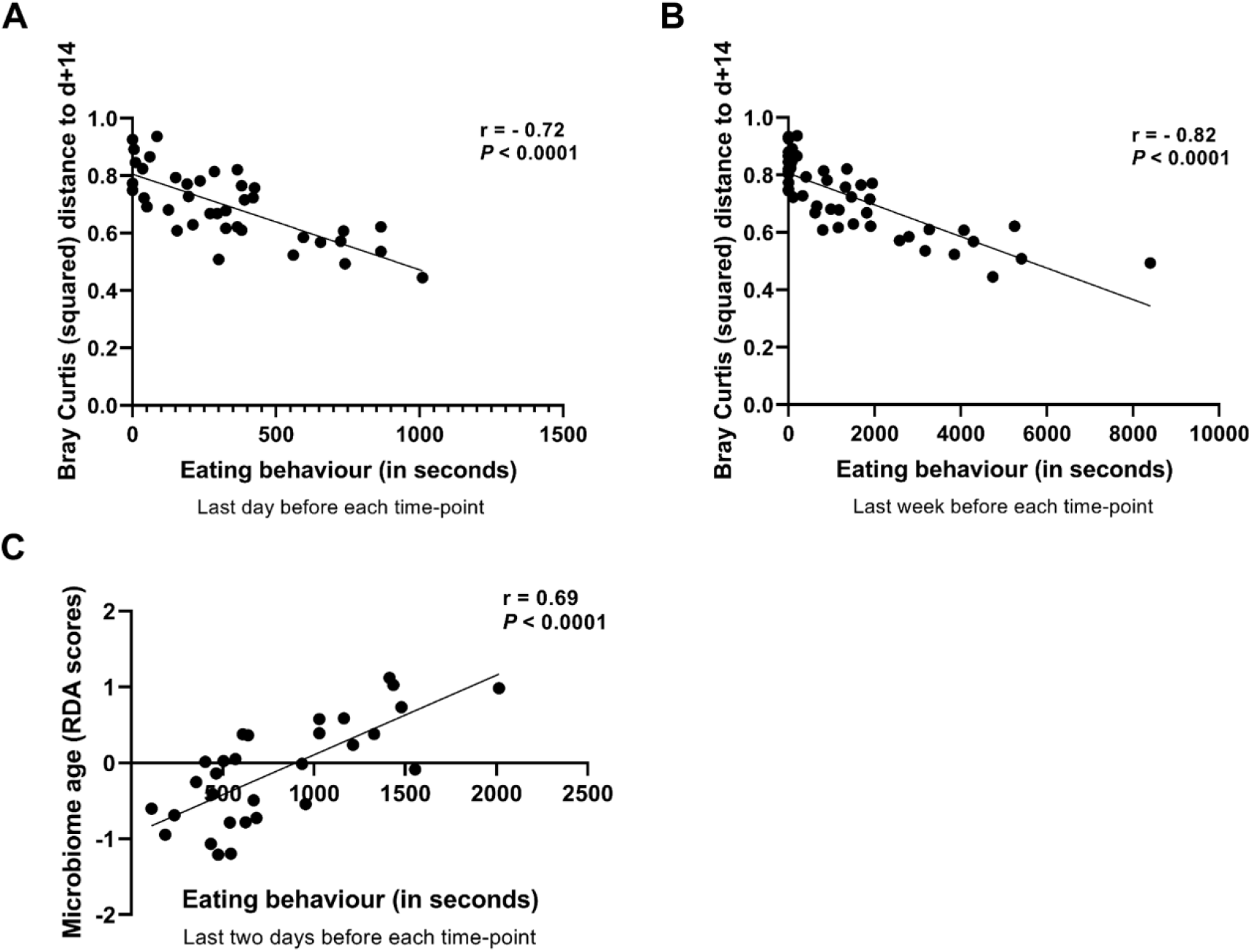
Spearman correlation between eating scores of individual piglets (n = 10) and squared Bray Curtis distance to their day+14 “matured” time-point: **(A)** last day eating score before each corresponding time-point (r = - 0.72, *P* < 0.0001) and **(B)** last week before each corresponding time-point (r = - 0.82, *P* < 0.0001). **(C)** Spearman correlation between eating scores (last two days from day15 timepoint) and their ‘microbiome age’ (r = - 0.69, *P* < 0.0001).

